# Unusual starch grains from the *Euphorbia caducifolia* Haines

**DOI:** 10.1101/2023.03.10.532015

**Authors:** Kusuma Venumadhav, Kottapalli Seshagirirao

## Abstract

Nature has chosen the starch as a universal form for storing carbohydrate. In granule form, it is semi-crystalline, water-insoluble, dense and a ubiquitous constituent. Structure of the starch granules depends on their biological source and varies from part to part of the sources. In sequence to that, starch granules from the latex of *Euphorbia caducifolia* was isolated and characterized under different conditions to reveal its uniqueness in the structure. A flat dumbbell-shaped starch was notified with distinctive “maltese cross” under various microscopes. These starches are found to be high amylose contents, which resist water solubility, swelling index, and turbidity of the starches. X-Ray Diffraction studies states the presence C type pattern of crystallinity whereas typical polysaccharide spectrum was observed with ATR studies. Gelatinization of the starches remarked them as highly thermal stable and retrogradation studies failed to detect realignment of the heat disrupted amylose and amylopectin. Lintnerization of starch granules reveals the high complexity of the structure. Thus, latex starch granules from the *E caducifolia* are considered to be unusual because of their highly complex structure, shape and properties.

## 1. Introduction

Starch is a major carbohydrate reserve in plants and stored in tubers and seeds of the plants as granules. The structure and shape of an individual starch granule is strictly under genetic and metabolic control, thus the granules isolated from different parts of the same plant may differ morphologically (French, 1984). It’s known that, amylose and amylopectin are the constituents of the starch granules and their orientation, content ratios decides physical parameters like solubility, swelling nature etc.

Granule structure of the starch under genetic control, therefore the starch granules from different parts of the same plant may differ in their structure. Some non-articulated milky laticifers of Euphorbiaceae contain unusual rod to spindle, osteoid to dumb-bell and discoid shaped starch grains. Previously, characteristic shapes of Euphorbiaceae latex starch granules were reported by Rafn (1798), Groom (1889), Molisch (1901), Solereder (1908).. Further, Czaja has reported structure and classification of the starch granules in vascular plants (Czaja, 1969, 1978). Later, seshagiriao and Prasad reported typology of latex starch granules and their importance in systematics(Seshagirirao & Prasad, 2001, 1988) Subsequently, reports on latex starch granules have been dropped because of their complexity and identifying the sources.

The present investigation aimed to study the structure and properties of latex starch granules from *Euphorbia caducifolia* to reinforces the starch granules of various Euphorbia species.

## 2. Materials and Methods

### 2.1 Isolation and Purification

The latex of *Euphorbia caducifolia* was collected and frozen in liquid nitrogen, frozen latex was thawed and centrifuged at 20,000 x g for 1 hour at 4°C. The supernatant was removed and sediments were dissolved in hexane to remove the fatty acids, oils and etc. The sediments remained were dried at room temperature. The dried powder was washed with citrate buffer pH 6.0 for 2-3 times and suspended in water. From the suspension, the starch grains were isolated by centrifugation in 2M sucrose at 2000 X g for 30 min. the sediment starch grains were washed with 0.2 M phosphate buffer pH 7.2 and distilled water for 3 times (Seshagirirao & Prasad, 1988). The prepared starch grains were used for microscopic studies and Physiochemical properties.

### 2.2 Microscopic studies

Latex starch grains were studies with a light microscope. Tincture iodine was used as a stain. Scanning Electron Microscope studies were done by transferring the grains to metallic stubs, dried in a desiccator. The starch grains were coated with Palladium-gold (Pd/Au) and observed under SEM at 15-20 kV(Mahlberg, 1973). The confocal laser scanning microscope was used to analyze structural architecture. A Leica 60x/1.4NA/oil immersion was used to study the grain structure. Ar^+^ laser was used as the excitation source and by using a set of PMTs, photoluminescence images were recorded (Manca et al., 2015). 2% Starch grains were prepared in distilled water containing 0.02% sodium azide as a preservative. Further, observed thinly sectioned starch granules under Transmission Electronic Microscope (Mamoru, Keiji, & Dexter, 1979). For sectioning, Starch granules were prefixed in fixative composed with glutaraldehyde (3%) along with sucrose (0.1 M) in 50 mM sodium cacodylate buffer at pH 7.5 and incubated in osmium tetraoxide (2.5%) for 2.5 h. After the post-fixation, granules were dehydrated with successive percentages of ethanol (50%, 70%, 80% and finally 100%). Dehydrated specimens were washed with acetone 3 times and embedded in resin and sectioned using the ultratome. Sectioned specimens were observed under TEM.

### 2.3 Blue value (BV) and amylose content

The blue value of the complex (Iodine-starch) was measured at 680 nm. In brief, starch was dissolved in acetate buffer pH 4.8 and iodine and potassium iodide added to it and measured the blue value (BV) (Takeda, Takeda, & Hizukuri, 1986).

### 2.4 Solubility, Swelling Index and Turbidity

Starch granules were analyzed for its solubility in water along with swelling index according to a method of Li & Yeh (2001). The starch granules (db) was added to the water I a screwcapped tube and vortexed for 10 s. Incubated for 30 min at temperatures 50, 60, 70, 80 and 90°C with additional mixing at 2 min interval. Then, cooled to room temperature and centrifuged for 30 min at 2000 g. The cloudy layer and sediments were separated and dried to constant weight. The weight of supernatant (WL) and sediments (WS) was used to calculate the water solubility Index (WSI) and Swelling Index (SI) as follows (Kong et al., 2016):

WSI= WL/0.1 ×100%

SI= WS/[0.1 × (100% - WSI)] (gg^-1^)

Turbidity was measured as mentioned by Lan et al. (2008). The water suspension of starch was boiled for 1 hrs with constant stirring and cooled to 30°C. The turbidity of the solution was determined with Chemito UV2100 spectrophotometer at 620 nm against a blank (distilled water) (Reis, Cristiana, & Beirão-da-costa, 2012)

### 2.5 Infra Red spectroscopy analysis

The starch granules for analyzed for the functional groups referred to the standard potato starch. FT-IR analysis was performed by using Platinum– Alpha, a single reflection Attenuated Total Reflection(ATR) module. The spectrum of the starch granules was measured from 4000 cm^-1^ to 500 cm^-1^.

### 2.6 X-ray diffraction Analysis

Starch granules are semi-crystalline in nature. The crystalline behavior of the granules was studied using X-ray powder diffractometer. The diffractogram of the starch was measured in *2*^®^ between 5 and 40° with 1° /min scan rate.

### 2.6 Thermal and retrogradation properties

Differential scanning calorimeter (DSC) used to study the thermal characteristics of the starch. Gelatinization of the starch was investigated under the natural condition and different water proportions. The pan was sealed after addition of water and equilibrated at room temperature for 2 hrs. The experiment was performed from 30 to 120°C with heating rate 5°C/min and the empty sealed pan was used as a reference. After cooled down, the pan was kept at 4 °C and −20°C for 1 week and rescanned for retrogradation properties.

### 2.8 Lintnerization

The starch granules were subjected to mild acid hydrolysis with 2.5 N HCl for 5 days at 30°C and aliquots of the treated samples at different time points was diluted and centrifuged. The supernatants were analyzed for sugars released at various temperatures and time points (Bertoft, 2004).

## 3. Results and Discussions

### 3.1 Microscopic analysis

Starch granules from the Euphorbia latex are diverge in shape and structure in comparison to other starch grains like rice, maize, corn and other. The characteristic starch grains were isolated from latex and analyzed under various microscopes. Dumbbell-shaped starch was noticed with a light microscope and confirmed with iodine staining (Fig. 1). Scanning electron microscope revealed its structure with the size of 40-60 μm where maximum size of potato starch grains is 110 μm (Fig. 2). Previously, similar type of starches have been reported from *Euphorbia cattimandoo, E. milli, E. nerifolia, E. tirucalli* and also from its other family members like *Pedilanthus tithymaloides* and *Synadenium grantii* (Seshagirirao & Prasad, 1988). Latex starch granules are grow in apposition and granules at early stage are round or ovoid in shape but on mature, they stretch to characteristic shapes like discoid, rod or dumb bell (French, 1984) (Fig. 3). Molecular orientation of the starch architecture gives the birefringent pattern under polarized light. Dark Maltese cross was reported with avocado starch in 1987 by Kahn (Kahn, 1987) and in potato starch (French, 1984).

**Fig. 1.**
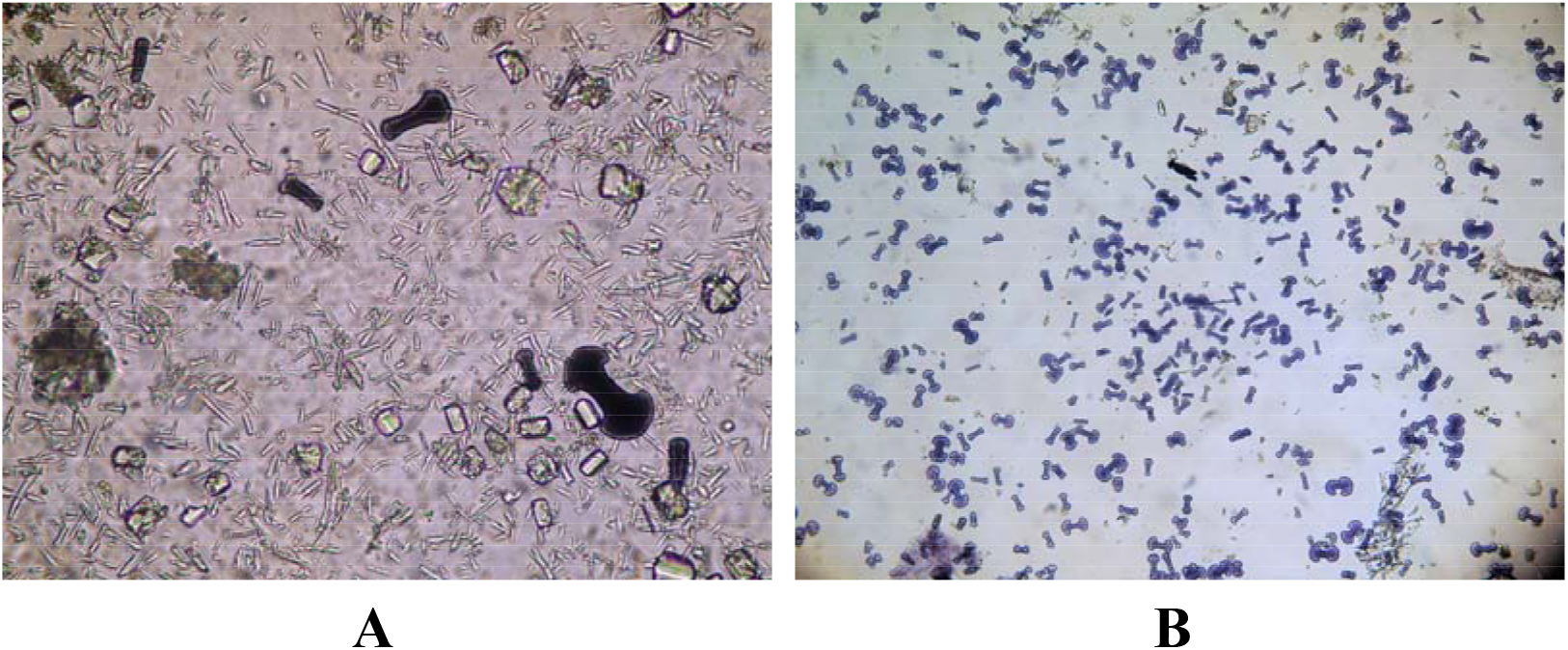
Iodine starch granules from *Euphorbia caducifolia* latex A) Starch granules in the latex along with calcium oxalate crystals, B) Purified starch.

**Fig. 2.**
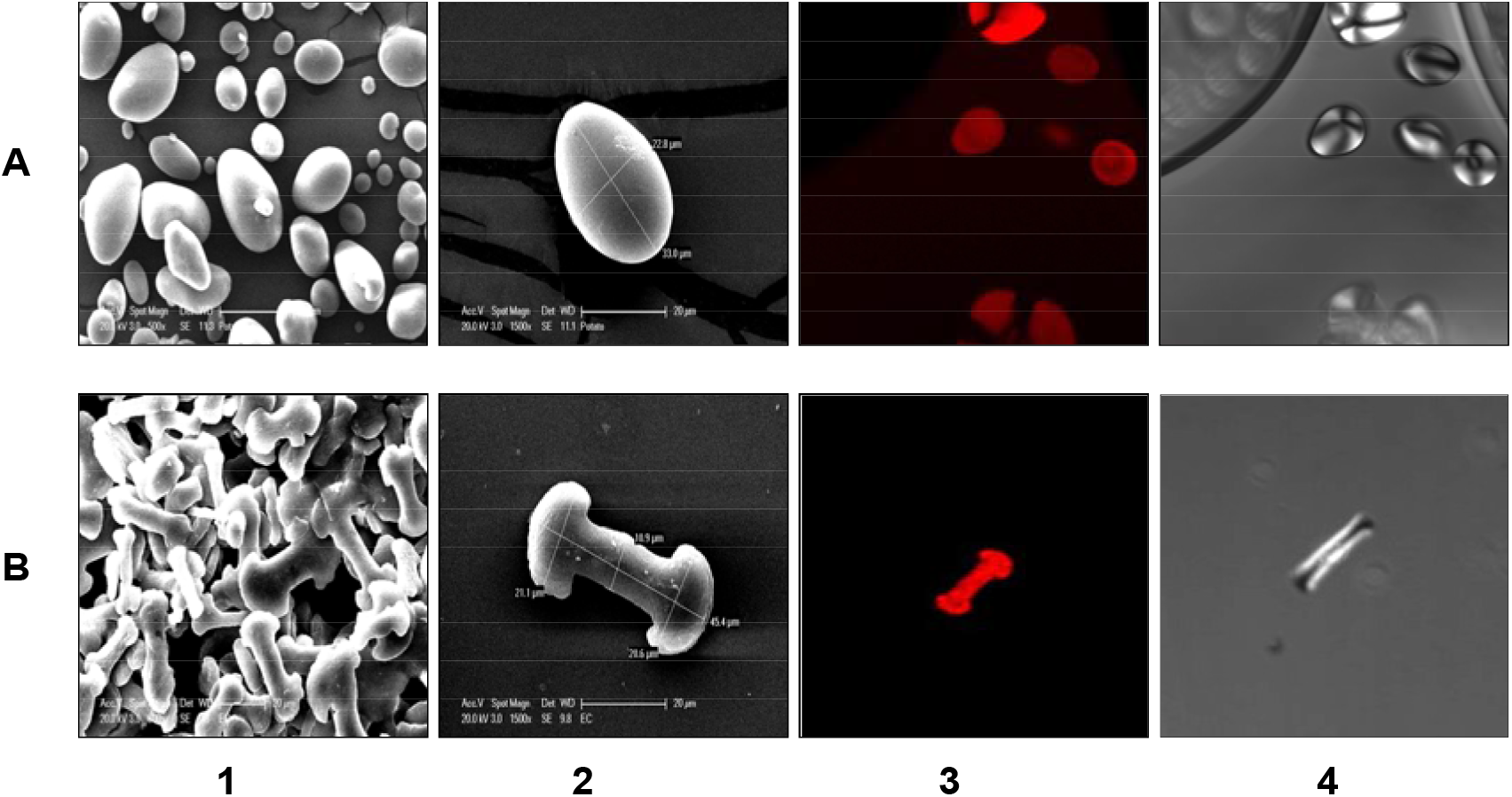
Microscopic view of Starch granules. A) Potato Starch, B) *Euphorbia caducifolia* Starch; 1,2-Scanning Electron microscope; 3, Confocal Laser microscopy, 4, Polarized light.

**Fig. 3.**
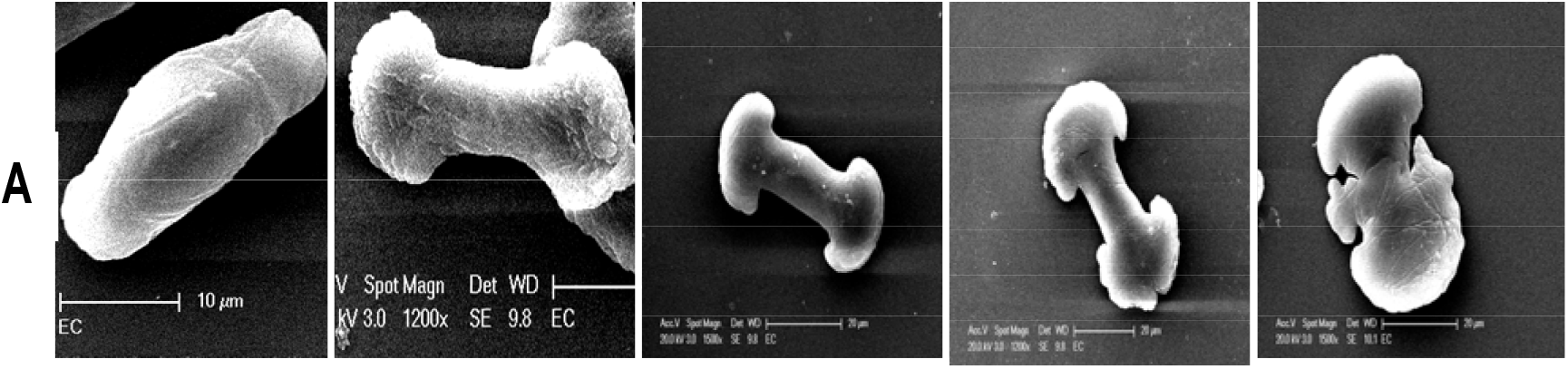
Structural View of starch granules. A) The growth process of starch granules (Left to Right) young to mature stages.

Confocal laser microscopy divulges the internal structure of the starch granules. Generally, starch granules are made up of amylose and amylopectin from the midpoint of the granule termed as “Hilum”. At lower levels of the structure, the alternate arrangement of amorphous and crystalline regions are made due to the higher level arrangement by clustering of side chains branching of radially arranged amylopectin. This appearance of the periodicity of starch granules is a universal feature and irrespective of botanical sources (Daniel J Gallant, Bouchet, & Baldwin, 1997). The almost identical molecular architecture was noticed with *E. caducifolia* starch (Fig. 4). Alternative arrangements or growth rings formed were noted and the same was confirmed by TEM studies. TEM studies spotted the growth rings with 100-400 nm thick and smooth edges of starch (Fig. 5). It can observe central cavity of the starch granule which is universal.

**Fig. 4.**
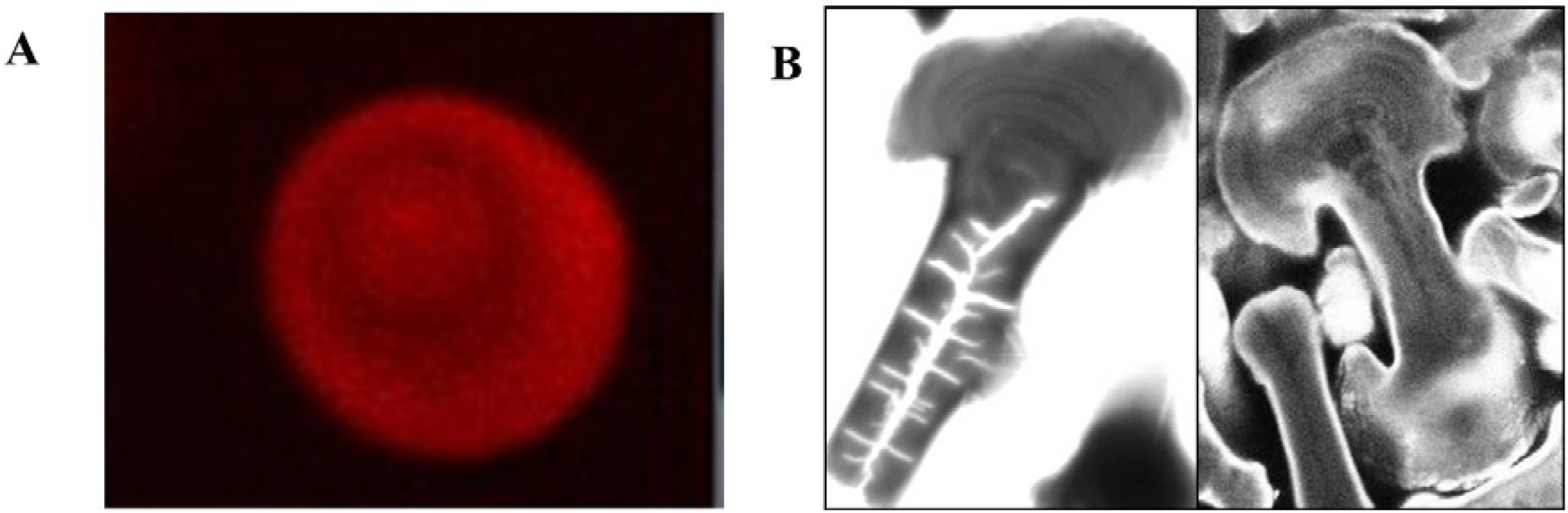
Starch granule Architecture A) Potato starch; B) *Euphorbia caducifolia* starch;

**Fig. 5.**
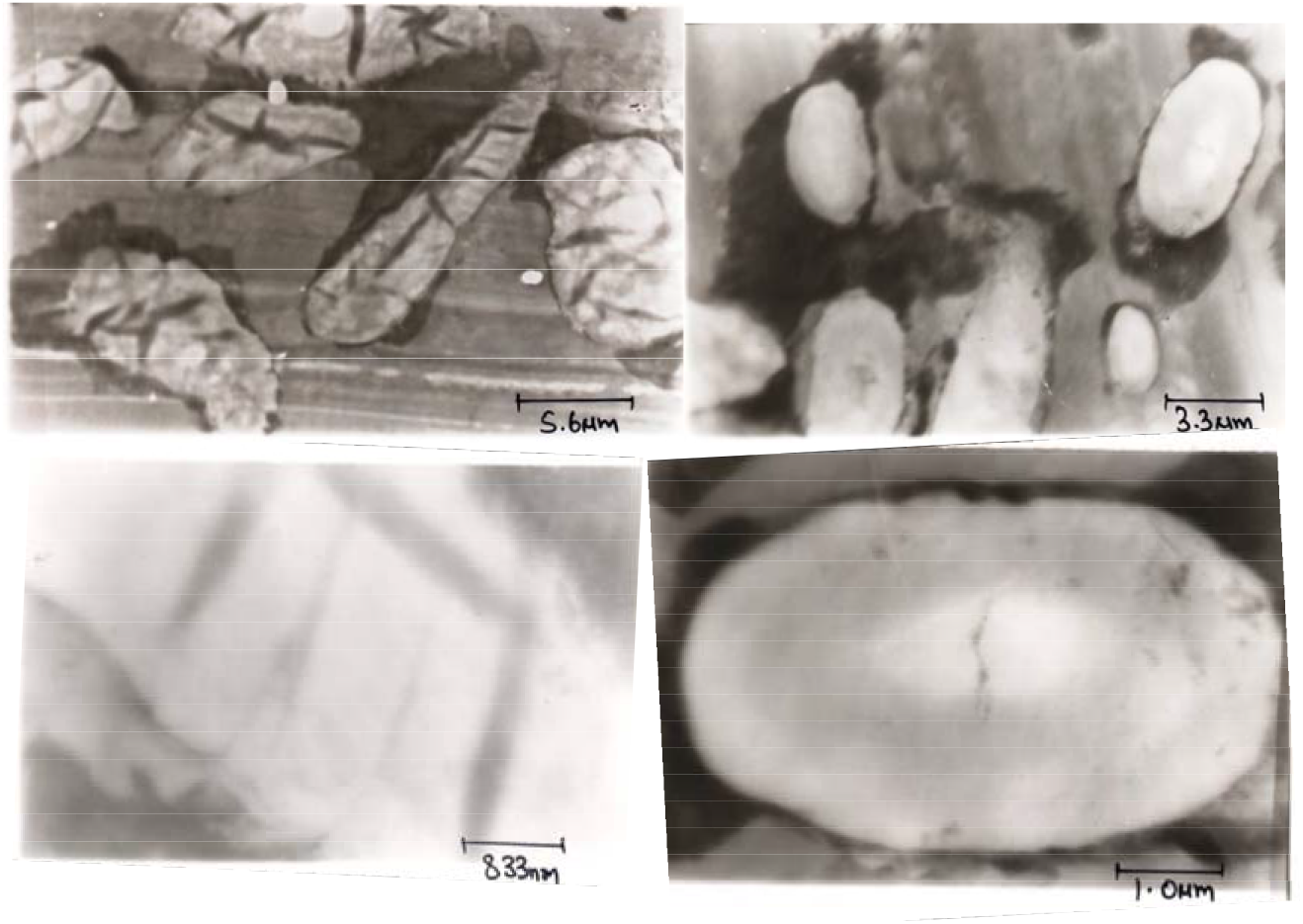
Transmission Electron Microscopy of sectioned starch granule.

### 3.2 Physicochemical properties and Amylose contents

Amylose contents were found to be 45% where normal starch grains from rice, wheat, and maize were found to be 20-33%. This implies the high contents of the amylose presents in EC starch. Further, the presence of high content amylose in EC starch was supported by high blue values with Iodine staining in comparison with potato (Fig. 6). Chemically, Iodine forms a blue colour complex with amylose of the starch and red to purple colour complex with amylopectin (Colin & Claubry, 1814; Foster et al., 1943; Stromeyer, 1815). These coloured complexes results suggest the structures possessing intermediate between amylopectin and amylose. Absorption studies of iodine and starch complex at 680 nm implies the greater contents of amylose were presents in EC starches than the potato starches.

**Fig. 6.**
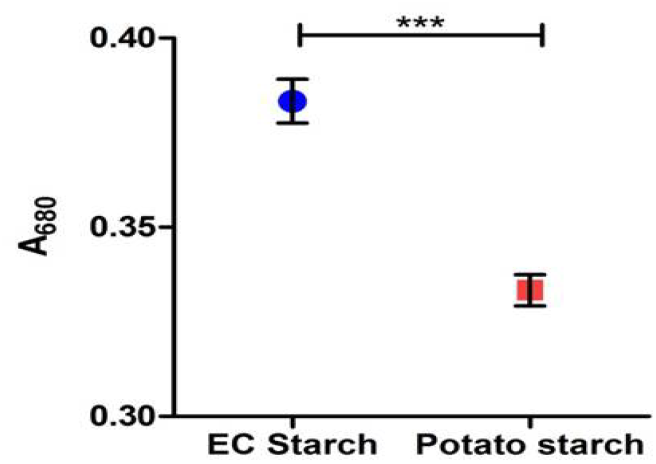
Blue value of Iodine starch complex.

Physicochemical properties like solubility, swelling index and turbidity were analyzed in comparison with reference potato starch (Fig. 7). The swelling index decreased and not completely solubilized even after 100°C with EC starch whereas potato starch completely solubilized above 80°C. This results in the complexity of the molecular structure of EC starch granule. The starch granules with high amylose contents swell to a low degree since starch granules are more rigid and resistant to digestibility (Sandhya Rani & Bhattacharaya, 1989). EC starch shows lower turbidity value than potato starch. Lower turbidity due to reluctant nature settled starch granules (Lan et al., 2008).

**Fig. 7.**
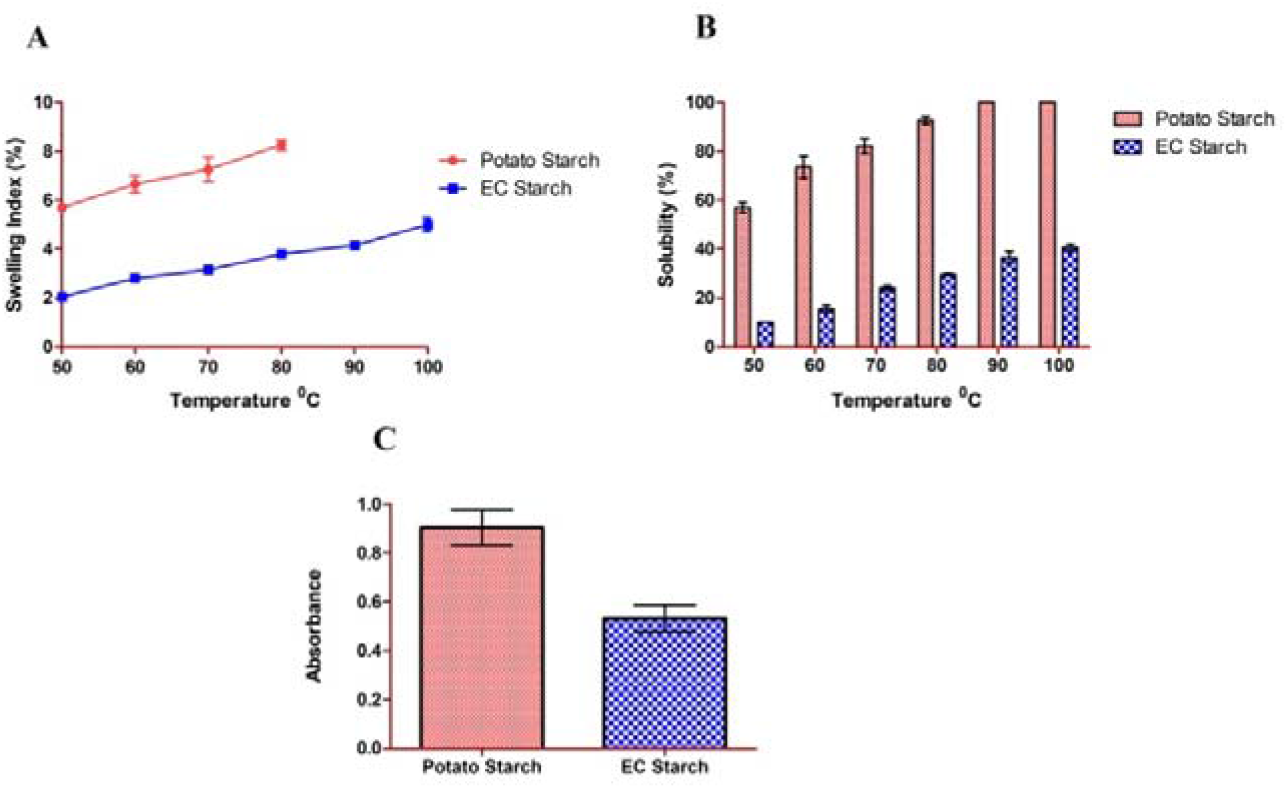
Physicochemical properties of starches; A) Swelling index, B) Solubility, C) Turbidity.

### 3.3 Crystallinity and functional group analysis

Starch granules isolated were analyzed for crystallinity and variations in the functional group present. Crystal structure released the presence of the intermediate composition of Amylose and amylopectin. EC starch shows the C-type crystallinity whereas potato with A-type. Starch have 15-45% of crystallinity varies botanical sources. Thus, starch granules display “Maltese cross” under polarized light (D J Gallant et al., 1992; Zobel, 1988). It has been known that starch granules from the EC and other Euphorbiaceae are devoid of “Maltese cross” (Spilatro & Mehlberg, 1985). Therefore, the crystallinity of the starch granules exhibits intermediate i.e C-type crystallinity. XRD patterns suggest the C-type crystallinity of the EC starch (Fig. 8) with peaks at 13°, 15°, 23° and 25°. Infra-Red spectra of EC starch and potato starch are look similar which suggest the absence of impurities.

**Fig. 8.**
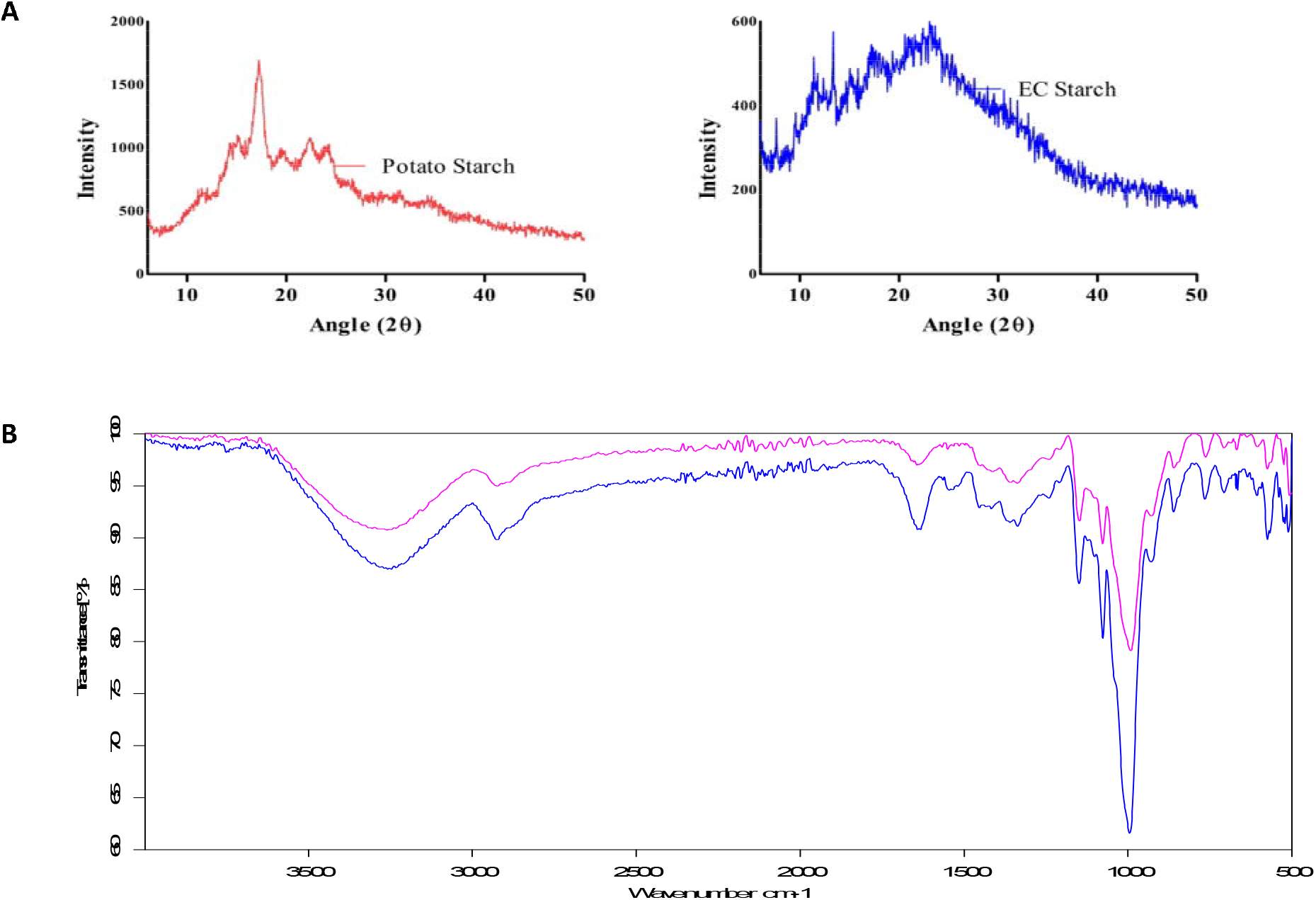
A) X-ray diffractions of potato and EC starch; B) IR spectrum of the starches potato (pink), EC starch (blue)

### 3.4 Thermal and retrogradation properties

Differential Scanning Calorimetry was used to study the gelatinization properties of starch granules (Table 5.1). When a molecule heated, thus results in the loss of weight and absorption or release of the energy occurs as a result of phase transition. Starch granules with 5% water and 5% NaCl result in the endothermic peaks at 58.89°C and 61.22°C respectively. Change in the enthalpy was 24 and 21 J. Gelatinization temperatures and change in the enthalpy closely resembles the corn starch (Jane et al., 1999). The apparent amylose content of starch correlated to onset and end set of temperatures. High amylose content lowers the onset temperature. EC starch granules do not exhibit any retrogradation properties with 5% water. (Wang et al., 2015).

**Table 5.1.** Thermal properties of EC starch.

| Sample | Onset | Thermal | | $\Delta H$ |
| --- | --- | --- | --- | --- |
|  |  | End set | Peak |  |
| **EC Starch** | **0** | **0** | **0** | **0** |
| **EC Starch + 5% Water** | **50.47** | **69.41** | **58.89** | **24.176** |
| **EC Starch + 5% NaCl** | **52.77** | **70.54** | **61.22** | **21.593** |

### 3.5 Lintnerization

Lintnerization is the process of acid treating starch. Acid catalyzes the starch was reported by Nageli in 1874 (Nageli, 1874). Acid digests the amorphous regions of the starch granules and left crystalline regions are called lintners.This has been first reported by Lintner in 1886 (Lintner, 1886). Lintnerization reduces the molecular weight of the starch. Lintnerization was observed with potato as well as with EC starch.The rate of lintnerization is high for the EC starch in comparison to potato starch. EC starch lintners are constant after 10 days with 20% of lintners whereas potato starch shown after 7 days with 5 %. Later, the lintners at every stage were noted under Scanning Electron Microscope (Fig. 9). Effect of lintnerization on various starch granules shows the complexity of the starches (Jayakody & Hoover, 2002).

**Fig. 9.**
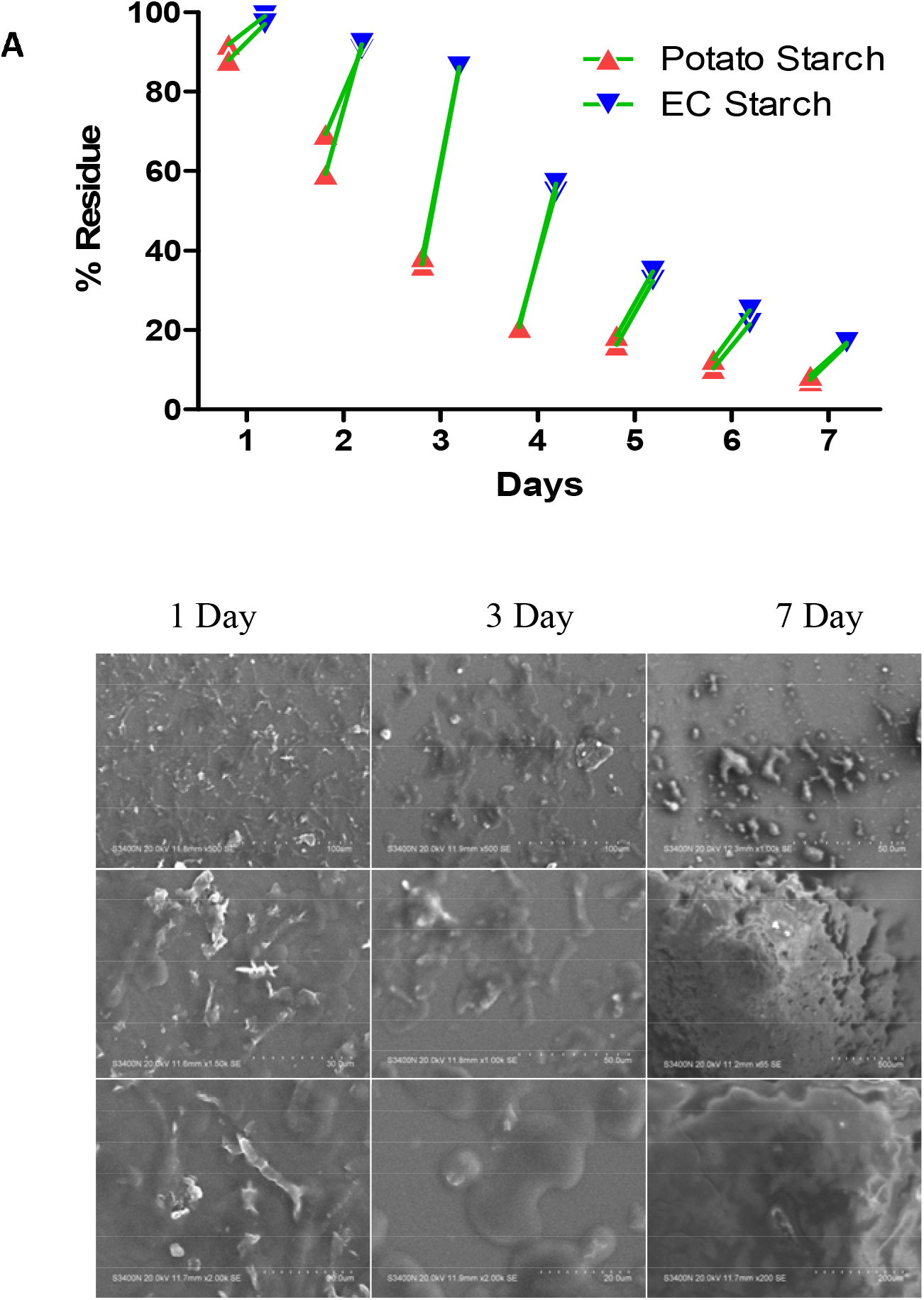
Lintnerization of starch A) Percentage of lintners, B) Lintner images on SEM.

## Conclusions

Starch granules from *E. caducifolia* latex are flat dumb bell shape. Laser Confocal microscope revealed the birefractive nature and growth rings of the starch granules as like potato starch. Granules grow from round or ovoid (young) to full mature dumb bell or discoid shapes. Blue values of the starch disclose the high contents (45%) of amylose with C-type crystallinity. High amylose presence decreased the solubility, swelling ability, turbidity, and lintnerization. Typical starch spectrum was identified with IR spectra with devoid of retrogradation properties.

